# Human iPSC-derived Olfactory Sensory Organoids model olfactory function and drug response

**DOI:** 10.64898/2026.11.24.769047

**Authors:** Vanshita Singh, Aishwarya Pantula, Kaustubh Joshi, Om Sadul, Nitya Lagadapati, Ashis Kumar Srikumar, Naigang Li, Xiaolei Zhu, Vasiliki Mahairaki, Annie Kathuria

**Author notes:** Funding: Xiaolei Zhu (R56MH135895, R01HL180044).

## Abstract

Intranasal drug delivery is being explored as an alternative approach to overcome the blood-brain barrier in the treatment of central nervous system diseases. However, little is known about how therapeutics administered through this route affect human olfactory tissue. Here, we develop human induced pluripotent stem cell-derived olfactory sensory organoids that reproduce key cellular and functional features of the developing olfactory system. Over three months of differentiation, the organoids acquire diverse neuronal and glial-associated populations, including cells expressing markers of olfactory sensory neurons and interneurons. Single-nuclei RNA sequencing shows changes in cellular composition and lineage-associated transcriptional programs during maturation, while comparison with a human fetal nasal atlas identifies shared neural populations and developmental signatures. Alongside these molecular changes, the organoids develop spontaneous electrical activity and coordinated network bursting. Exposure to the glioma-relevant kinase inhibitors Dasatinib, Trametinib, and Selumetinib produces distinct changes in neuronal network activity, demonstrating the ability of the model to capture compound-associated functional responses. Together, these findings establish a human Olfactory Sensory Organoid system for investigating olfactory development and evaluating how therapeutics may affect olfactory neural tissue during intranasal drug administration.

**ToC figure:** 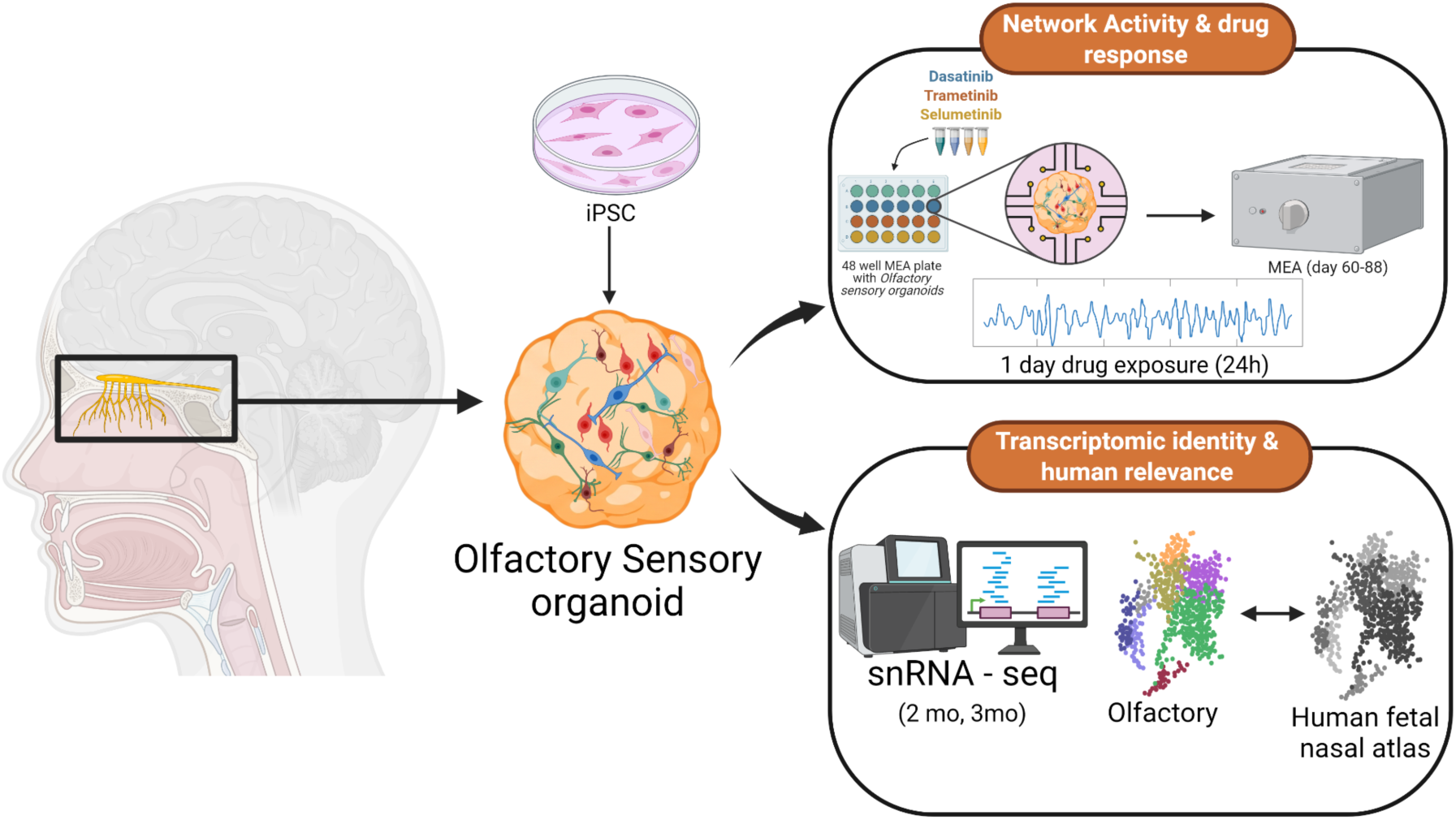

## 1. Introduction

Glioblastoma (GBM) is the most common malignant primary brain tumor in adults and accounts for approximately half of all malignant primary central nervous system (CNS) tumors. ^[1]^ Despite maximal surgical resection followed by radiotherapy and chemotherapy, median overall survival remains approximately 15 months.^[2]^ Diffuse infiltration, extensive intratumoral heterogeneity (ITH), and treatment resistance contribute to poor therapeutic outcomes. ^[3]^ Molecularly targeted therapies have also shown limited clinical efficacy, in part because achieving adequate drug exposure within the CNS remains challenging. ^[4]^ The blood-brain barrier (BBB) restricts the penetration of many systemically administered therapeutics into the brain. ^[5]^ Although BBB integrity is disrupted in portions of GBM, it can remain relatively intact at infiltrative margins, contributing to heterogeneous drug exposure across the tumor and surrounding brain tissue. ^[6,7]^ These limitations have motivated alternative CNS delivery strategies, including convection-enhanced delivery and intranasal administration, that reduce dependence on systemic transport across the BBB. ^[8,9]^

Intranasal administration offers a potential route for CNS drug. ^[10,11]^ The olfactory epithelium contains olfactory sensory neurons (OSNs), sustentacular cells, basal progenitors, and other supporting populations. ^[11]^ OSNs extend dendrites into the nasal cavity and project axons through the cribriform plate to the olfactory bulb, establishing a direct anatomical interface between the peripheral olfactory system and the brain. ^[12]^ Molecules administered intranasally can access CNS compartments along olfactory and trigeminal pathways, although transport efficiency depends on molecular properties, formulation, nasal deposition, and clearance. ^[9,10]^ This route has therefore been investigated for compounds with limited BBB permeability, including candidate therapeutics for brain tumors. ^[13]^ However, the effects of such therapeutics on human olfactory neural tissue, particularly during repeated or prolonged exposure, remain poorly characterized.

Investigation of human olfactory tissue has been limited by the difficulty of obtaining and maintaining primary olfactory neuronal populations in vitro. Animal models, particularly rodents, have therefore provided much of our understanding of olfactory development, sensory transduction, neuronal regeneration, and circuit organization. ^[14]^ However, substantial molecular differences distinguish the human and rodent olfactory systems. Humans possess approximately 400 functional olfactory receptor (OR) genes, compared with approximately 1,000 in mice. ^[15]^ A substantially larger fraction of the human OR repertoire consists of pseudogenes. ^[16] [17]^ Functional properties can also diverge between orthologous mammalian ORs, with differences in ligand sensitivity, efficacy, and odorant recognition. ^[18][19]^ These differences in receptor repertoire and function complicate direct extrapolation of olfactory responses from rodents to humans and support the development of complementary human experimental models.

Human pluripotent stem cells offer an alternative approach for generating olfactory lineage cells and investigating human olfactory development and function. Directed differentiation protocols combining dual-SMAD inhibition with developmental morphogens, including fibroblast growth factor 8 (FGF8), have enabled the generation of olfactory lineage populations from human pluripotent stem cells.^[20–22]^ More recently, human induced pluripotent stem cells (iPSCs) have been differentiated into OSNs exhibiting molecular and functional responses to odorant. ^[23]^

Human olfactory-derived cells have also been used for pharmacological studies. Cultured olfactory epithelial-derived neuronal cells have shown responses to dopaminergic, serotonergic, and glutamatergic stimulation. ^[24]^ Olfactory epithelial biopsies and neurosphere-derived cells have been used to assess pharmacodynamic responses and drug-induced changes in cell behavior. ^[25][26]^ More recently, studies have demonstrated functional responses in mature human OSNs, including odorant- and IBMX-evoked activity in acute olfactory epithelial tissue and selective odorant responses in iPSC-derived OSNs. ^[23,27]^ Pharmacological rescue of axonal injury has also been demonstrated in iPSC-derived OSNs, establishing the feasibility of therapeutic intervention in this system. ^[28]^ However, these studies have largely relied on isolated neuronal populations, progenitor cultures, or acute tissue preparations rather than multicellular three-dimensional models capable of reporting coordinated network-level responses.

Three-dimensional organoid cultures extend these approaches by supporting the development and interaction of multiple neural cell populations within the same system. ^[29,30]^ This added cellular complexity may better capture the coordinated maturation and network behavior that cannot be assessed in isolated neuronal populations or acute tissue preparations. However, whether iPSC-derived olfactory models develop both the cellular diversity and functional network activity required for pharmacological testing remains unclear.

Single-cell and spatial transcriptomic studies of adult and developing human olfactory tissue have established molecular references for basal progenitors, immature and mature OSNs, and other olfactory-associated populations. ^[11,31]^ single-nuclei RNA (snRNA) sequencing can therefore be used to resolve heterogeneous neuronal and glial populations in vitro and evaluate their correspondence with developing human tissue. Mature OSNs express canonical odorant-transduction components and typically select a single olfactory receptor from a large genomic repertoire. ^[13,32]^ Multielectrode arrays (MEAs) further complement transcriptomic profiling by enabling longitudinal measurements of spontaneous spiking and network bursting in living cultures as a functional assay. ^[33]^

Here, we generated human iPSC-derived olfactory sensory organoids (OSO) through staged neural induction, FGF8-mediated olfactory specification, and long-term maturation. We characterized their cellular and molecular development across three months using immunofluorescence, snRNA sequencing, and pseudotime analysis, and benchmarked their transcriptional identity against a developing human fetal nasal atlas through shared clustering and directional concordance analysis. We used longitudinal multielectrode array recordings to assess the emergence and maturation of spontaneous neuronal network activity. Followed by an evaluation of the functional responses of maturing OSOs to the glioma-relevant kinase inhibitors Dasatinib, Trametinib, and Selumetinib. Together, our analyses establish a human OSO model that integrates olfactory lineage identity, developmental state, and network-level function. We hope our system provides a human-specific framework for studying olfactory neurobiology and for evaluating the functional effects of therapeutics that may encounter olfactory neural tissue during intranasal CNS delivery.

## 2. Results

### 2.1. FGF8-guided differentiation generates structurally mature olfactory sensory organoids

We developed a staged differentiation strategy to generate OSOs from human iPSCs (**Figure 1A,B**). We first aggregated iPSCs into embryoid bodies (EBs), followed by dual-SMAD inhibition from days 2-8 to establish a neural lineage. We then withdrew the dual-SMAD inhibitors and applied FGF8 from days 8-12 to specify olfactory fate. At day 12, we transferred the EBs to 6 weeks continued maturation. We treated cultures with DAPT from days 30-60 to promote neuronal differentiation and subsequently maintained the OSOs in N2/B27 medium through functional characterization and drug treatment.

**Figure 1.**
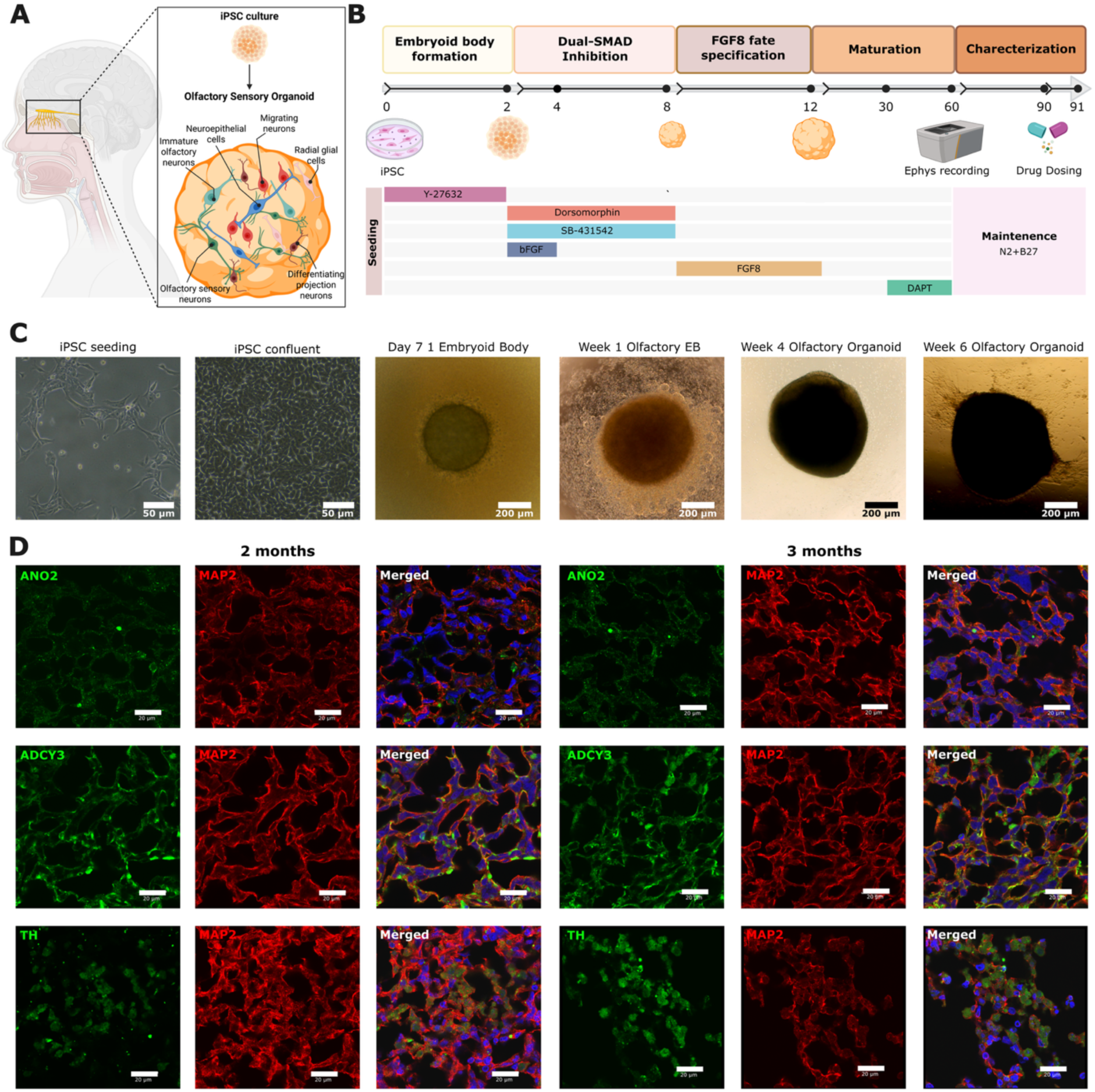
Generation and phenotypic characterization of human iPSC-derived OSOs. (**A**) Schematic overview of the iPSC derived OSO model. (**B**). Differentiation timeline for OSO generation. EB formation (days 0-2), dual-SMAD inhibition with dorsomorphin and SB-431542 (days 2-8), and FGF8-mediated olfactory specification (days 8-12), followed by adherent maturation. We treated OSOs with DAPT from days 30-60 and maintained them in N2/B27 medium through electrophysiological characterization and drug dosing at ∼day 88. (**C**) Representative brightfield and phase-contrast images showing morphological progression from iPSC seeding and confluent iPSC cultures through EB formation and OSO maturation. Images show a day-7 EB, a week 1 olfactory EB, a week 4 OSO, and a week 6 OSO with extensive peripheral neurite outgrowth. Scale bars: 50 µm for iPSC images and 200 µm for EB and OSO images. (**D**) Representative immunofluorescence images of 2- and three month OSOs stained for ANO2, ADCY3, or tyrosine hydroxylase (TH) (*green*), together with the neuronal marker MAP2 (*red*) and nuclear counterstain DAPI (*blue*). ANO2 and ADCY3 were detected within MAP2-positive neuronal regions at both developmental stages, while TH labeled a more spatially restricted neuronal population. Scale bars: 20 µm. Schematic in **(A**) and (**B**) created in BioRender. Kathuria, A. (2026) https://BioRender.com/8rqyz3g.

The cultures underwent progressive morphological maturation across the differentiation period (**Figure 1C**). Compact EBs formed within the first week and reached approximately 350-450 µm in diameter by day 7. During early olfactory specification, EBs increased to approximately 450-500 µm and retained a smooth, optically dense morphology. By week 4, adherent OSOs reached approximately 550-600 µm in diameter. Following DAPT (a Notch inhibitor) treatment, OSOs developed prominent peripheral neurite outgrowth, which became readily apparent by week 6, while the central OSO body remained approximately 600-700 µm in diameter, ready for functional assessment.

### 2.2. Olfactory sensory organoids acquire sensory neuronal and interneuronal features

We next evaluated olfactory neuronal identity in two and three month OSOs by immunostaining for ANO2, ADCY3, TH, and the neuronal cytoskeletal marker MAP2 (**Figure 1D**; **SI Figure 1; SI Table 1, Supporting Information**). MAP2-positive neuronal regions were distributed throughout the OSO sections at both developmental stages, consistent with the extensive neurite formation observed during adherent maturation. ^[34]^

We detected ANO2 within MAP2-positive neuronal regions at both two and three months. ANO2 encodes the calcium-activated chloride channel TMEM16B, which contributes to amplification of the olfactory sensory transduction response.^[35]^ At two months, ANO2 staining appeared relatively sparse and localized to discrete neuronal profiles. ANO2 remained detectable at three months, where positive structures were distributed across the MAP2-positive neuronal network. This persistence supported maintenance of an olfactory sensory neuron-associated phenotype during extended culture.

ADCY3 showed a broader pattern of expression than ANO2 and was readily detected within MAP2-positive neuronal regions at both developmental stages. ADCY3 is a ciliary adenylyl cyclase that links odorant receptor activation to intracellular cAMP signaling and represents a central component of canonical olfactory sensory transduction.^[36]^ The presence of both ADCY3 and ANO2 therefore showed that the OSOs expressed multiple components of the olfactory transduction machinery rather than neuronal markers alone. ADCY3 staining remained prominent at three months, indicating preservation of this sensory-associated molecular program during OSO maturation.

TH-positive cells displayed a more restricted distribution than either ANO2 or ADCY3. At two months, TH-positive cells were dispersed within MAP2-positive neuronal regions, whereas by three months the TH-positive signal appeared more spatially clustered within defined neuronal populations. TH catalyzes the rate-limiting step in dopamine synthesis and is characteristic of dopaminergic interneuron populations within the olfactory bulb.^[37]^ The distinct TH staining pattern therefore supported the presence of an olfactory-associated interneuron-like population in addition to the broader sensory neuronal compartment.

Together, the persistence of ANO2, ADCY3, and TH at both developmental stages supported maintenance of multiple olfactory-associated neuronal phenotypes, while differences in their spatial distributions suggested continued organization of these populations between two and three months. We subsequently examined these populations using snRNA sequencing and pseudotime analyses to determine whether the apparent age-dependent organization was accompanied by changes in cellular or developmental.

### 2.3. snRNA transcriptomic profiling reveals diverse cell populations in human olfactory sensory organoids

To characterize the cellular composition of OSOs during differentiation, we performed snRNA sequencing at two and three months (**Figure 2A**). Following quality control and filtering, we retained 1,720 nuclei from two-month OSOs and 1,565 nuclei from three-month OSOs for downstream analysis. We show the quality control of these nuclei, including UMIs per nucleus, genes per nucleus, and mitochondrial content in **SI Figure 2** (**Supporting Information**). We used independent Leiden clustering to identify 14 transcriptionally distinct clusters at two months and 12 clusters at three months.

**Figure 2.**
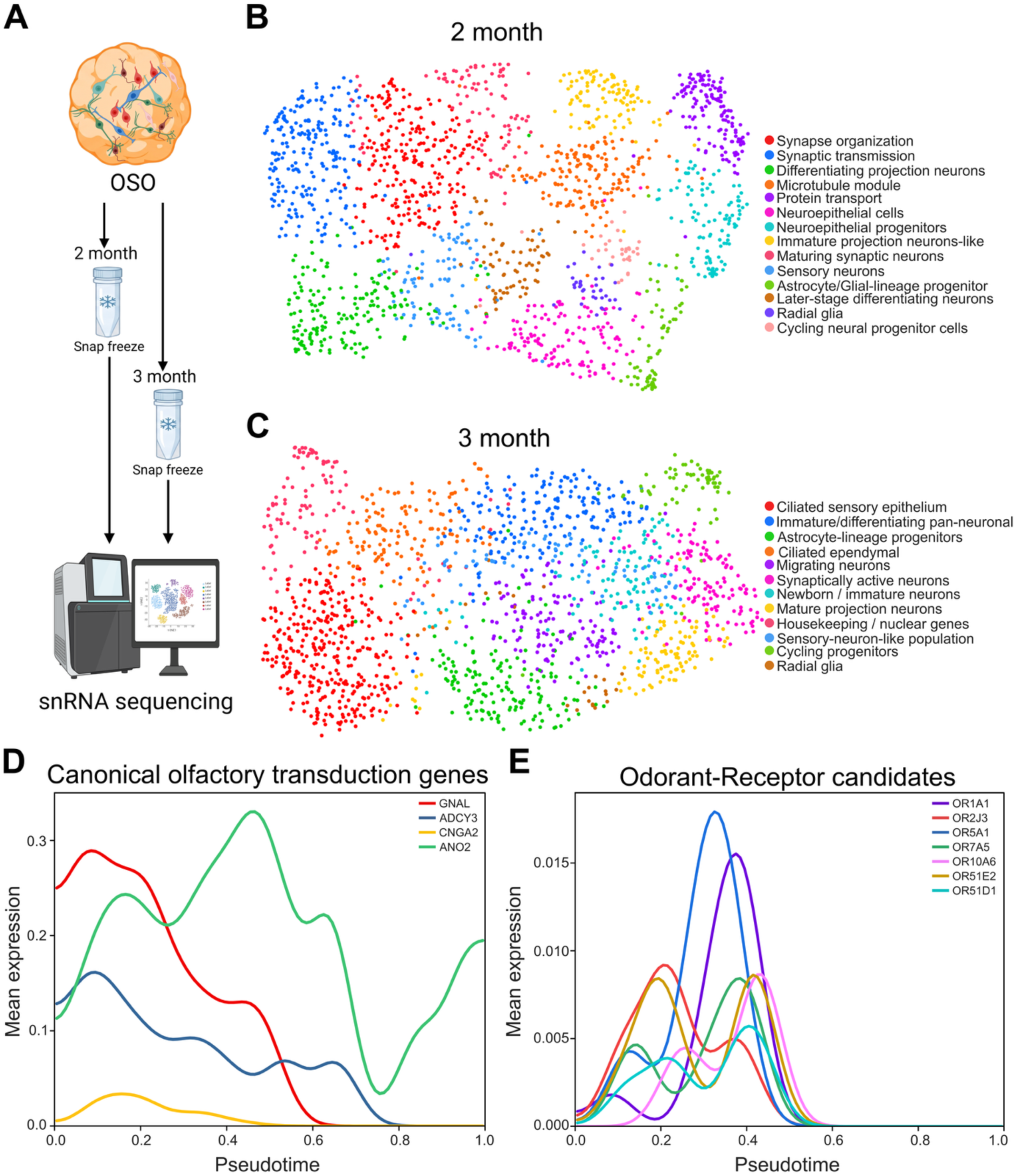
snRNA sequencing of OSO at two and three months. (**A**) OSOs were snap-frozen at two and three months for snRNA sequencing. (**B**) UMAP of 1,720 nuclei from two-month OSOs, colored by 14 Leiden clusters. Progenitors were 18.4% of nuclei, differentiating and immature projection neuron-like populations 16.6%, synaptic and later-differentiating populations 37.6%, sensory neurons 5.3%, and the glial-lineage progenitor 4.3%. (**C**) UMAP of 1,565 nuclei from three-month OSOs, colored by 12 Leiden clusters. Ciliated sensory epithelium was the largest cluster (21.5%; 26.8% with sensory-neuron-like nuclei). Mature projection neurons were 6.7% and astrocyte-lineage progenitors 9.9%. Immature, migrating, newborn, and synaptically active neurons together were 35.5%. Cycling progenitors (4.2%) and radial glia (1.5%) remained. (**D**) Mean expression of the canonical olfactory transduction genes GNAL, ADCY3, CNGA2, and ANO2 plotted against pseudotime for the three-month OSO (**E**) Mean expression of the odorant-receptor candidates OR1A1, OR2J3, OR5A1, OR7A5, OR10A6, OR51E2, and OR51D1 plotted against the same pseudotime axis. Schematic in (**A**) Created in BioRender. Kathuria, A. (2026) https://BioRender.com/8rqyz3g.

At two months, the OSOs contained heterogeneous populations of progenitors and differentiating neurons (**Figure 2B**). Neuroepithelial cells, neuroepithelial progenitors, radial glia, and cycling progenitors collectively accounted for 18.4% of nuclei, while differentiating and immature projection neuron-like populations represented 16.6%. Synaptic and later-differentiating neuronal populations constituted the largest group (37.6%), whereas sensory neuron-like cells and glial-lineage progenitors accounted for 5.3% and 4.3%, respectively. At three months, the OSOs exhibited a greater proportion of cells expressing sensory-associated transcriptional signatures (**Figure 2C**). Ciliated sensory epithelial cells represented the largest individual cluster (21.5%), and together with sensory neuron-like populations, accounted for 26.8% of nuclei. Mature projection neurons and astrocyte-lineage progenitors represented 6.7% and 9.9%, respectively. Cycling progenitors (4.2%) and radial glia (1.5%) remained detectable, while immature, migrating, newborn, and synaptically active neuronal populations collectively accounted for 35.5% of nuclei. These findings revealed differences in the relative abundance of progenitor, neuronal, and sensory-associated cell populations between the two differentiation stages.

To examine olfactory sensory neuron-associated transcriptional programs, we analyzed the expression of canonical olfactory transduction genes along the inferred pseudotime trajectory of three-month OSOs (**Figure 2D**). GNAL and ADCY3 exhibited their highest expression during early pseudotime, followed by a gradual decline. In contrast, ANO2 expression increased through intermediate pseudotime and remained elevated toward the later stages of the trajectory. CNGA2 expression remained comparatively low throughout pseudotime, with a maximum expression approximately tenfold lower than that of GNAL. We further examined the expression of seven odorant receptor genes, OR1A1, OR2J3, OR5A1, OR7A5, OR10A6, OR51E2, and OR51D1 (**Figure 2E**). At the individual nucleus level, at least one of the seven receptor transcripts was detected at 2.4%. Together, these findings demonstrated the presence of olfactory sensory-associated transcriptional programs and detectable odorant receptor expression in a subset of OSO nuclei, although expression of individual olfactory transduction components remained heterogeneous.

### 2.4. Pseudotime analysis identifies distinct sensory, interneuron, and glial transcriptional programs

To further characterize the transcriptional programs associated with olfactory lineage differentiation, we performed diffusion pseudotime analysis independently on the two- and three-month snRNA sequencing datasets. We examined the expression dynamics of genes associated with mature olfactory sensory neurons, dopaminergic interneurons, GABAergic interneurons, and olfactory ensheathing glia (**Figure 3**).

**Figure 3.**
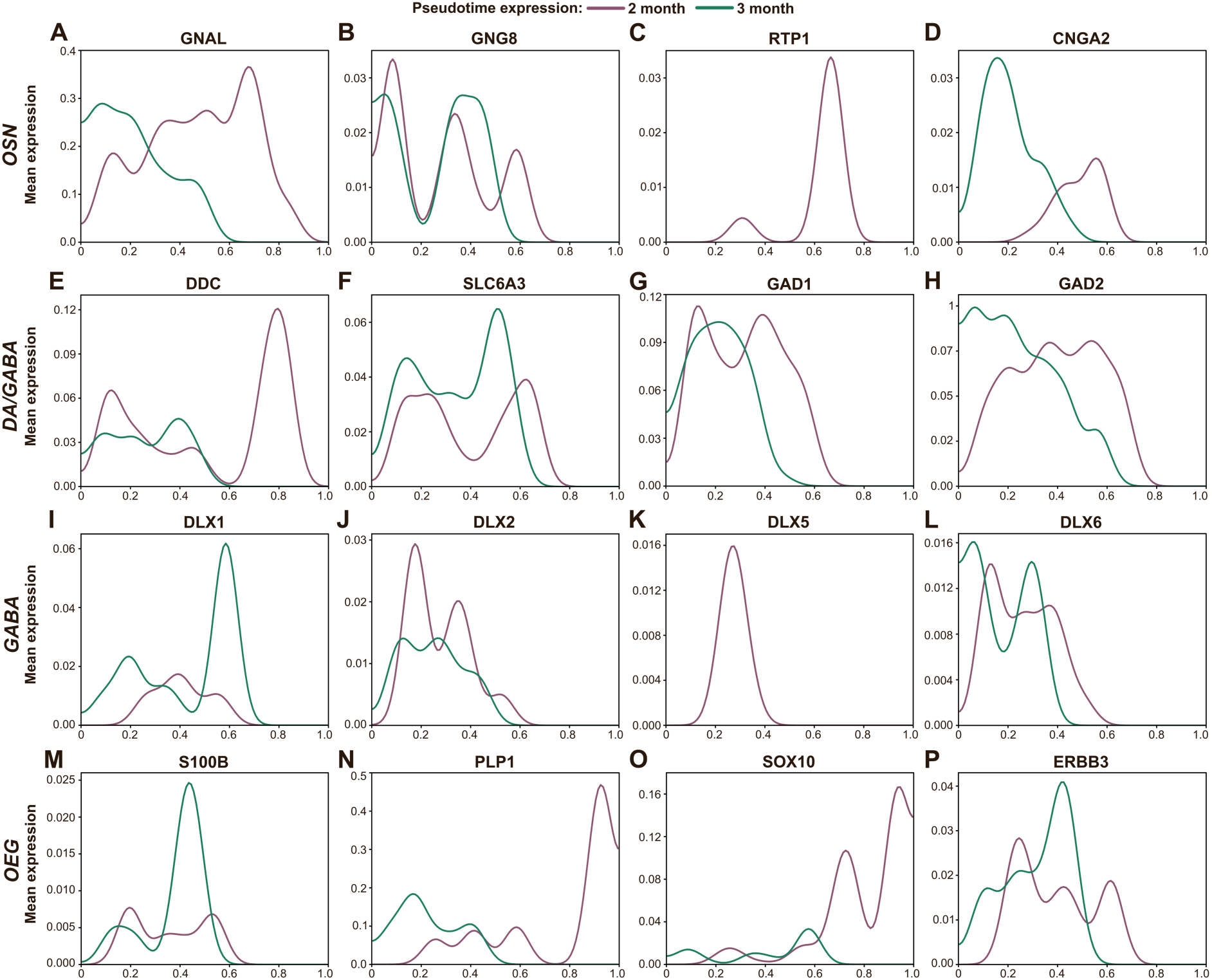
Mean expression along pseudotime in OSOs at two and three months. Each panel is one gene. *Purple* is the two month OSO and *green* is the three month OSO, plotted on the same axes. We ordered each age on its own pseudotime, shown from 0 to 1, and the y axis is mean expression, scaled separately for each gene. **(A-D)** OSN development genes: GNAL, GNG8, RTP1, and CNGA2. **(E, F)** Dopaminergic/GABAergic (DA/GABA) genes: DDC and SLC6A3. **(G-L)** GABAergic genes: GAD1, GAD2, DLX1, DLX2, DLX5, and DLX6. **(M-P)** OEG developmental genes: S100B, PLP1, SOX10, and ERBB3.

Genes associated with mature OSNs displayed distinct pseudotemporal expression patterns at the two differentiation stages (**Figure 3A-D**). At two months, sensory-associated markers were distributed across the trajectory, with GNAL and RTP1 reaching their highest expression toward later pseudotime and CNGA2 peaking within the intermediate-to-late region. At three months, GNAL, GNG8, and CNGA2 expression was concentrated toward earlier pseudotime and declined along the trajectory, whereas RTP1 expression remained minimal.

We next examined genes associated with periglomerular dopaminergic interneurons (**Figure 3E,F**). At two months, DDC and SLC6A3 reached their highest expression toward the end of the trajectory. In contrast, both genes exhibited earlier expression peaks in the three-month dataset, with DDC showing a lower maximum expression than at two months.

GABAergic interneuron and granule-cell markers also exhibited distinct pseudotemporal distributions (**Figure 3G-L**). At two months, GAD1, GAD2, DLX1, DLX2, DLX5, and DLX6 showed relatively broad expression across early and intermediate pseudotime. At three months, GAD1, GAD2, and DLX6 were predominantly expressed toward early pseudotime, while DLX2 showed comparatively lower expression across early-to-intermediate pseudotime. In contrast, DLX1 exhibited a pronounced expression peak later in the trajectory, whereas DLX5 expression was largely restricted to the two-month dataset.

Finally, we investigated the expression dynamics of developmental genes associated with olfactory ensheathing glia (OEG) (**Figure 3M-P**). S100B and ERBB3 exhibited higher expression peaks at three months, particularly within intermediate pseudotime. In comparison, PLP1 and SOX10 reached their highest expression toward the end of the two-month trajectory. Collectively, these findings demonstrated distinct pseudotemporal distributions of sensory neuronal, interneuron, and glial-associated transcriptional programs between two and three months of differentiation.

Consistent with the age-dependent organization observed by immunostaining, pseudotime analysis showed redistribution of olfactory sensory, GABAergic/dopaminergic, and glial transcriptional programs between two and three months, supporting continued lineage remodeling during OSO maturation. ANO2 and ADCY3 remained detectable within MAP2-positive regions at both ages, and in the three-month trajectory ADCY3 was highest early while ANO2 increased through intermediate pseudotime and remained elevated. TH staining was dispersed at two months and more spatially clustered at three months, (**SI Figure 2, Supplementary information)** in parallel with DDC and SLC6A3, which peaked at the end of the two-month trajectory and earlier at three months .

### 2.5. Olfactory sensory organoids develop spontaneous network activity and respond to glioma-relevant kinase inhibitors

We next asked whether OSOs develop spontaneous, coordinated neuronal activity and whether pharmacological perturbation changes this activity (**Figure 4A**). Olfactory sensory neurons convert chemical stimuli into receptor potentials and action potentials, and spontaneous neuronal activity contributes to the development and maintenance of olfactory circuitry.^[27,38]^ We therefore used multielectrode array (MEA) recordings to complement the molecular and transcriptomic characterization of the OSOs and to determine whether coordinated network activity accompanies neuronal maturation. We recorded organoids weekly from day 60 through day 88. We then performed drug perturbation on day 88 and recorded network activity every 2 h for 24 h (**Figure 4A**).

**Figure 4.**
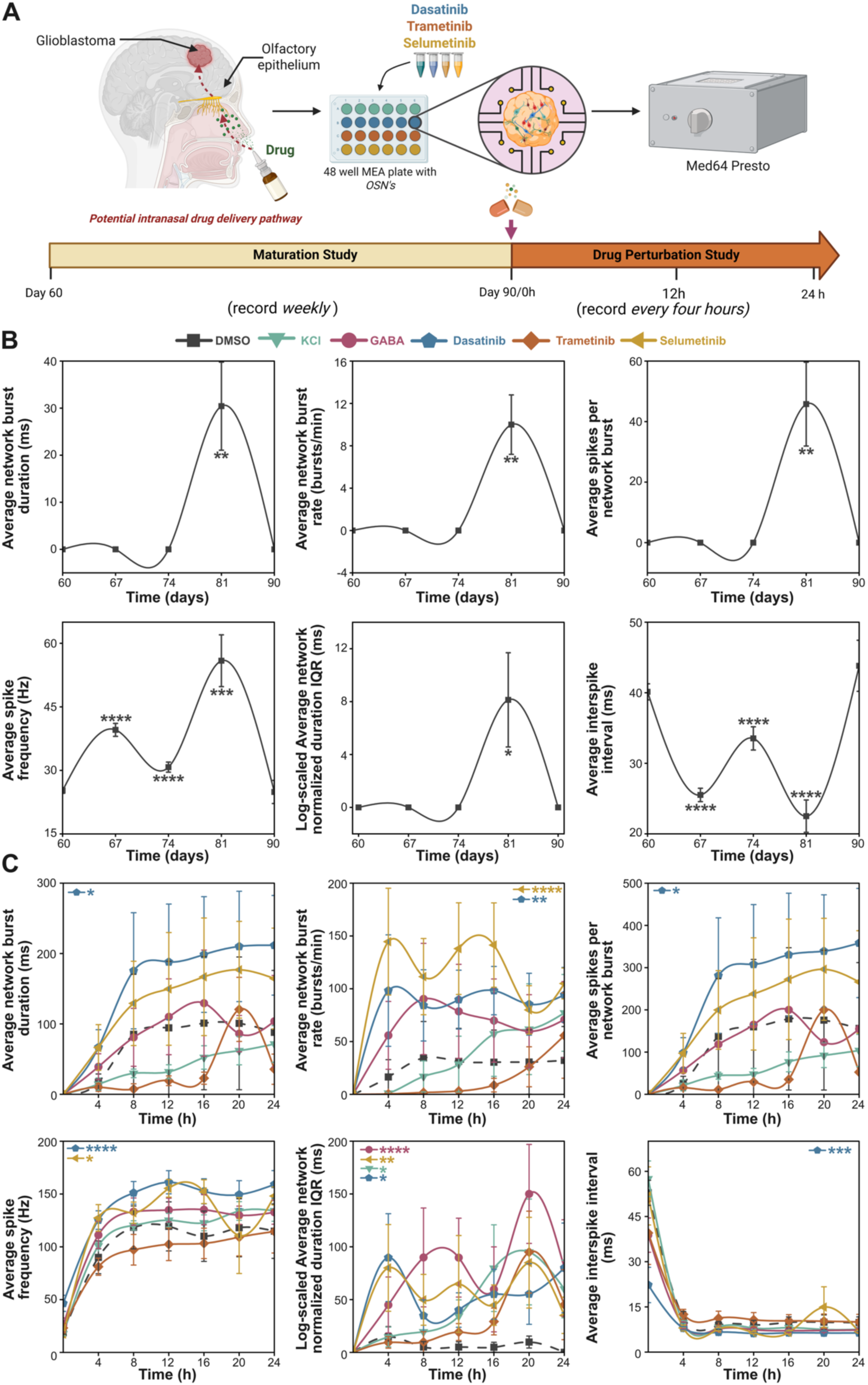
Electrophysiological maturation and drug-induced network responses in OSOs. (**A**) Experimental workflow for maturation (2 to 3 months) and drug perturbation studies on three month OSOs. We recorded spontaneous network activity from OSO weekly from day 60 (week 0) through day 88 (week 4) to assess functional maturation, (n=24). At approximately day 88, we treated mature OSO with DMSO vehicle, KCl, 25 µM, GABA 25 µM, 1 µM Dasatinib, 1 µM Trametinib, and 10 µM Selumetinib and recorded electrophysiological activity every 2 h for 24 h. (**B**) Maturation study. Average network burst duration, network burst rate, spikes per network burst, spike frequency, normalized network burst-duration IQR, and interspike interval. (**C**) Drug perturbation study for KCl, GABA, Dasatinib, Trametinib, and Selumetinib. For each treatment condition, we used OSO from three iPSC lines and averaged four OSO replicates per treatment. Data are presented as mean ± SEM. Dunnett’s multiple comparisons test against week 0 (**B**) or DMSO (C); *p < 0.05, **p < 0.01, ***p < 0.001, ****p < 0.0001; ns, not significant. Schematic in (**A**) created in BioRender. Kathuria, A. (2026) https://BioRender.com/8rqyz3g

Spontaneous electrical activity changed markedly across the maturation window (**Figure 4B**). Spike frequency rose from approximately 25 Hz at day 60 to 40 Hz at day 67 and peaked near day 81. It stayed significantly above day-60 levels at days 67, 74, and 81 (p < 0.0001, p < 0.0001, and p = 0.0002) and returned to baseline by day 88 (p = 0.9997). Coordinated network activity followed a similar trajectory. Network burst rate, spikes per network burst, and network burst duration stayed low in the earlier recordings and rose substantially around day 81. The interspike interval showed the opposite pattern. It fell significantly below day-60 levels at days 67, 74, and 81 (all p < 0.0001), reached its minimum near day 81, and returned to baseline by day 88 (p = 0.7201). These measurements place peak spontaneous and coordinated network activity at approximately 80 days of differentiation.

We then tested whether our three-month OSOs reported functional responses to kinase inhibitors relevant to glioma therapy. We selected Dasatinib, Trametinib, and Selumetinib as mechanistically distinct kinase inhibitors with established relevance to glioma signaling. Dasatinib targets Src-family kinase signaling and has been evaluated clinically in glioblastoma,^[4]^ whereas Trametinib and Selumetinib inhibit MEK within the MAPK pathway, a pathway investigators have pursued as a therapeutic vulnerability in glioma. ^[39,40]^ To benchmark these responses against canonical modulators of neuronal excitability, we included potassium chloride (KCl,25uM) as a depolarizing reference and gamma-aminobutyric acid (GABA, 25 µM) as an inhibitory reference alongside a DMSO treated control.

We recorded a 5-minute baseline without drugs, followed by treating the OSOs with each compound, and recording electrophysiological activity for 2 minutes every 2 hours for 24 hours (**Figure 4A,C)**. Dasatinib produced the broadest response and significantly altered all six network parameters relative to DMSO. It increased spike frequency (p < 0.0001), network burst rate (p = 0.0050), network burst duration (p = 0.0254), spikes per network burst (p = 0.0442), and normalized burst-duration IQR (p = 0.0104), and it lowered the interspike interval (p = 0.0003). These increases appeared rapidly and persisted across much of the recording period. Selumetinib also strongly increased coordinated activity, most prominently network burst rate (p < 0.0001), along with spike frequency (p = 0.0200) and normalized burst-duration IQR (p = 0.0091), although its temporal profile differed from that of Dasatinib. Trametinib produced a delayed response, with small early changes and increased burst-associated activity at later time points, but it did not differ significantly from DMSO on any parameter. KCl and GABA each significantly changed only normalized burst-duration IQR (p = 0.0437 and p < 0.0001). Dasatinib and Selumetinib therefore perturbed network activity more broadly than either reference modulator.

The interspike interval dropped sharply within the first 4 hours in the kinase-inhibitor conditions and stayed low for most of the 24-hour recording, although only Dasatinib lowered it significantly relative to DMSO. Trametinib and Selumetinib both inhibit MEK, yet they produced distinct electrophysiological profiles, and Dasatinib generated a separate pattern of sustained network activation. Together, these results showed that three month OSOs developed measurable spontaneous network activity and generate compound-specific functional responses to pharmacological perturbation

### 2.6. Olfactory sensory organoids align with neural populations of the human fetal nasal atlas

To benchmark OSO identity against developing human tissue, we integrated snRNA sequencing data from two-month and three-month OSOs with a published human fetal nasal atlas spanning post-conception weeks 7-12 (**Figure 5A**). ^[31]^ The integrated dataset contained 45,848 nuclei across the fetal atlas and both OSO ages, using 10,889 genes shared across datasets. Harmony integration resolved 24 shared clusters.

**Figure 5.**
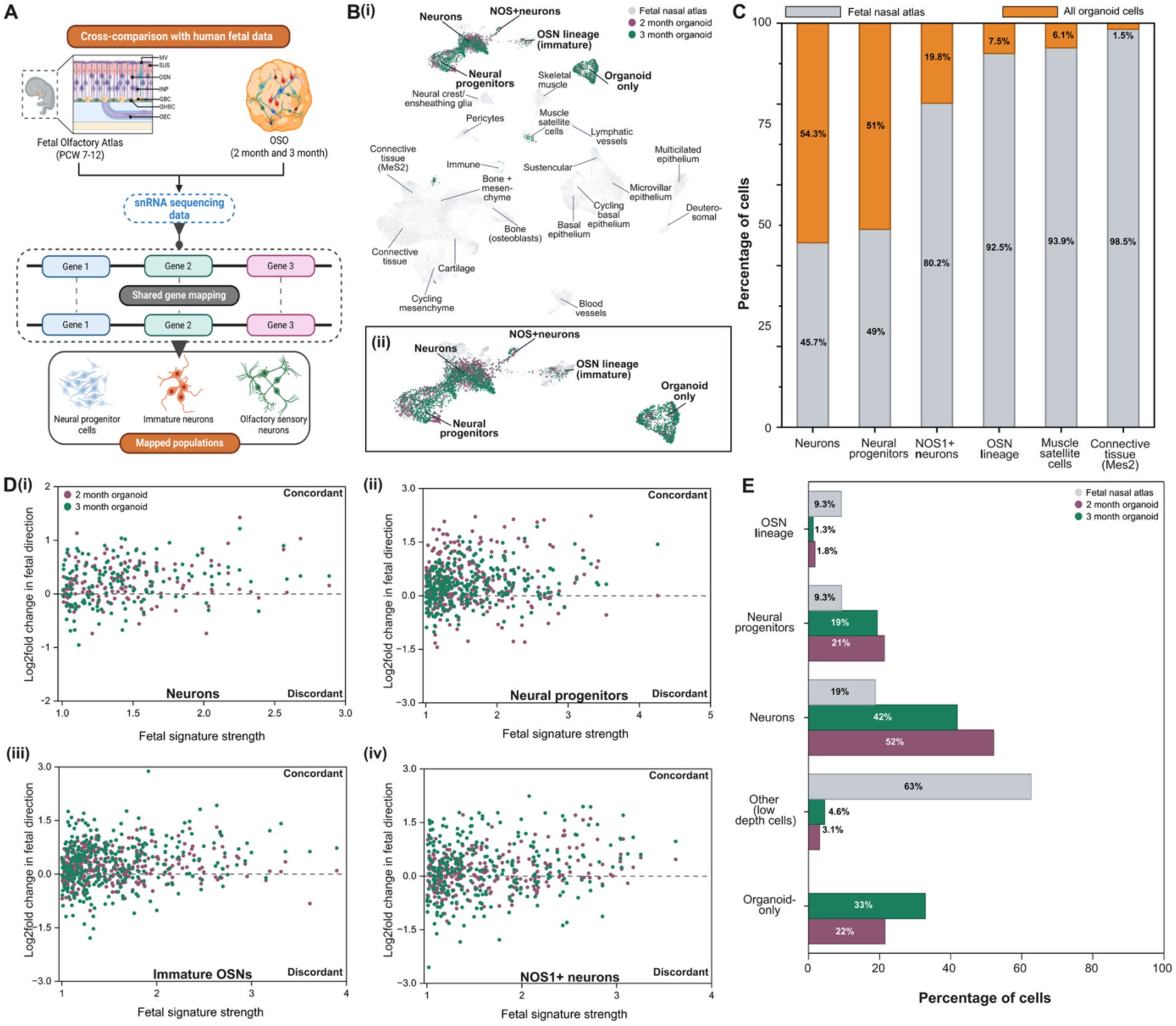
OSO nuclei align with neural populations of the human fetal nasal region. (**A**) Schematic of cross-comparison with human fetal data; Mbouamboua et al., 2026^[31]^ (**B**) Harmony-integrated UMAP of all 45,848 nuclei clustered by Louvain analysis into 24 shared populations. (**i**) Whole map and (**ii**) enlarged neural region. *Grey*, fetal atlas; mauve, two-month OSO; *green,* three-month OSO. Cluster labels reflect the fetal populations within each cluster; “organoid only” denotes a cluster containing no fetal nuclei. (**C**) Relative fetal and OSO contribution to shared clusters containing ≥1% OSO nuclei. OSO nuclei comprised 54.3% of the neuron cluster and 51.0% of the neural progenitor cluster, with lower representation in NOS1+ neurons, OSN lineage, muscle satellite cells, and connective tissue. (**D**) Directional concordance between fetal cluster signatures and corresponding OSO populations for (**i**) neurons, (**ii**) neural progenitors, (**iii**) immature OSNs, and (**iv**) NOS1+ neurons. Fetal signatures contained genes showing ≥2-fold enrichment or depletion relative to the other neural clusters and detected in ≥10% of nuclei on the higher-expression side. Values above zero indicate concordant direction between fetal and OSO populations; 50% concordance represents the chance expectation. Concordance at two and three months was 75% and 68% for neurons, 70% and 74% for neural progenitors, 68% and 67% for immature OSNs, and 57% and 61% for NOS1+ neurons, respectively. (**E**) Distribution of nuclei across major mapped populations within each sample. For the fetal comparison, percentages were calculated using only neuronal and olfactory epithelial nuclei from the reference atlas (n = 8,292). Neurons represented 52% and 42% of 2- and three month OSO nuclei, neural progenitors 21% and 19%, OSN lineage 1.8% and 1.3%, and the organoid-only population 22% and 33%, respectively. Schematic in (**A**) created in BioRender. Kathuria, A. (2026) https://BioRender.com/8rqyz3g.

OSO nuclei localized predominantly to the neural region of the integrated atlas (**Figure 5B i-ii**). Across both ages, 97% of OSO nuclei (3,871 of 3,973) mapped to clusters containing fetal neurons, neural progenitors, immature OSN lineage cells, NOS1+ neurons, or to a distinct organoid-only population. Both two-month and three-month OSOs contributed nuclei to each of these neural clusters. In contrast, OSO nuclei were largely absent from fetal mesenchymal, vascular, immune, epithelial, bone, cartilage, and glial populations. A small number of nuclei mapped to connective tissue, muscle satellite cell, and cycling mesenchymal clusters, but these nuclei had substantially lower gene detection and lacked the corresponding lineage markers.

We next quantified the relative fetal and OSO contribution to each shared cluster (**Figure 5C**). OSO nuclei comprised 54.3% of the neuron cluster and 51.0% of the neural progenitor cluster, indicating substantial overlap between OSO and fetal neural populations. OSO representation was lower in the NOS1+ neuron cluster (19.8%) and immature OSN lineage cluster (7.5%). Only minor fractions mapped to muscle satellite cells (6.1%) or connective tissue (1.5%). We ignored the organoid-only cluster. Counts for every shared cluster are listed in **SI Table 2** (S**upporting Information**). For integration with the human fetal nasal atlas, we used the quality-controlled datasets from AmpSeq, retaining 2,009 nuclei at two months and 1,964 nuclei at three months (**SI Figure 3, Supporting Information**).

We then tested whether transcriptional differences among OSO neural clusters followed the same direction as those observed among the corresponding fetal populations (**Figure 5D**). For each shared neural cluster, we defined fetal signature genes based on their relative enrichment or depletion compared with the other neural clusters and evaluated the direction of the corresponding change in two-month and three-month OSOs. Neuron signatures showed 75% and 68% concordance at two and three months, respectively, while neural progenitor signatures showed 70% and 74% concordance. Immature OSN signatures showed 68% and 67% concordance, whereas NOS1+ neuron signatures showed weaker agreement at 57% and 61%. Thus, the major OSO neural populations preserved the relative transcriptional organization of their fetal counterparts, although the magnitude of these differences was generally smaller in the OSOs. This may reflect the greater complexity of the in vivo fetal developmental environment, where diverse cellular interactions, spatially organized niches, and physiological signaling cues facilitate cellular maturation and transcriptional specialization to an extent not fully recapitulated in vitro.

Finally, we examined the distribution of nuclei within each sample independently (**Figure 5E**). Neurons represented the largest mapped population in both OSO ages, accounting for 52% of two-month and 42% of three-month nuclei, compared with 19% of the selected fetal neuronal and olfactory epithelial reference population. Neural progenitors accounted for 21% and 19% of 2- and three month OSO nuclei, respectively, while the OSN lineage represented 1.8% and 1.3%. The organoid-only population increased from 22% to 33% at three months. Together, these analyses showed that OSOs mapped predominantly to neural populations of the developing human nasal region, with strongest overlap with fetal neurons and neural progenitors and a smaller population aligning with the immature OSN lineage.

## 3. Discussion

Intranasal administration offers a potential route to deliver therapeutics to the CNS through pathways associated with the olfactory and trigeminal systems. However, the functional consequences of drug exposure on human olfactory neural tissue remain poorly understood. Existing human models, including isolated iPSC-derived OSNs, olfactory epithelial biopsies, neurosphere-derived cultures, and acute tissue preparations, have enabled cellular and pharmacological studies but provide limited access to coordinated responses across multicellular neural networks. In this study, we generated FGF8-directed differentiation human OSOs containing sensory neuronal, dopaminergic/GABAergic, and glial-associated populations, together with spontaneous network activity that matured over approximately three months. The coexistence of these populations and the emergence of coordinated electrical activity suggest that the organoids captured a level of multicellular organization not accessible in isolated olfactory neuronal cultures.

We observed age-dependent changes (two/three months) in olfactory sensory, GABAergic/dopaminergic, and glial-associated programs between two and three months, while pseudotime analysis showed redistribution across developmental trajectories rather than uniform progression along a single maturation axis. Olfactory receptor transcripts emerged in a subset of nuclei, further supporting progression toward OSN identity within a heterogeneous culture. Together, these findings indicate that prolonged culture supported continued developmental organization of the olfactory neural compartment rather than maintenance of a fixed differentiated state.

We next benchmarked the organoids against developing human olfactory tissue using the human fetal nasal atlas spanning PCW 7-12. ^[31]^ Approximately 97% of organoid nuclei localized within the neural region of the integrated transcriptional space, predominantly alongside fetal neurons, neural progenitors, immature OSN-lineage cells, and NOS1+ neurons. Neurons, neural progenitors, and immature OSN-lineage populations showed 67-75% directional concordance with corresponding fetal transcriptional signatures, whereas the smaller NOS1+ population showed more modest concordance. These findings suggest that the organoids preserved key relationships among developing human olfactory neural programs while retaining transcriptional features distinct from primary fetal tissue.

Longitudinal MEA recordings further showed that molecular maturation was accompanied by coordinated neuronal function. Several measures of network bursting and firing peaked near day 80 before declining in later recordings, suggesting functional reorganization over a simple progressive increase in activity. Spontaneous activity contributes to olfactory circuit development in vivo ^[38]^ , and neural organoid studies have similarly linked maturation with the emergence of organized spontaneous network activity. ^[33]^ We then tested three kinase inhibitors relevant to glioma biology: Dasatinib, which has undergone clinical evaluation in recurrent GBM ^[4]^ ; Trametinib, for which MEK1/2 inhibition has demonstrated antitumor and neuronal differentiation effects in GBM models ^[39]^ ; and Selumetinib, which has shown activity in molecularly defined NF1-mutant GBM models ^[41]^ . All three compounds altered network activity, and Trametinib and Selumetinib produced distinct temporal response profiles despite targeting the same kinase class. These findings indicate that shared molecular targets do not necessarily predict equivalent effects on human olfactory neural networks. In this study, we directly exposed olfactory neural tissue to defined drug concentrations and measured the resulting network-level responses. It therefore provides a controlled system for examining the functional response of human olfactory neural tissue after therapeutic exposure.

Beyond pharmacological testing, this system may also support studies of neurological disorders in which olfactory dysfunction emerges early. More than 95% of patients with Parkinson’s disease have been reported to exhibit olfactory impairment, which can precede motor symptoms^[42][43]^, while Alzheimer’s disease is similarly associated with substantial olfactory deficits.^[44,45]^ Neuropathological involvement of olfactory structures has been reported in both disorders, suggesting that patient-derived versions of this model could provide a human context for studying early disease-associated disruption of olfactory function. ^[46,47]^

Several limitations define the current scope of the model. The organoids primarily captured the neural component of the olfactory system and did not reproduce the complete sustentacular, epithelial, vascular, and immune compartments of the nasal mucosa. Incorporating epithelial support populations or adapting the system to an air-liquid interface could allow future studies to link drug transit with subsequent neural responses. We also tested a single concentration of each compound over 24 hours; dose-response studies, repeated dosing, and longer exposures will be required to determine whether the observed electrophysiological responses persist, adapt, or progress toward toxicity. Finally, independently analyzed patient-derived and genetically diverse iPSC lines will be important for defining donor-specific variability and extending the model toward disease-specific applications. Together, our findings establish a human OSOs model that integrates developmental identity, correspondence with primary human tissue, spontaneous network activity, and pharmacological responsiveness.

## 4. Experimental Section

### iPSC culture, thawing, maintenance, and cryopreservation

After thawing our iPSCs , we supplemented the culture medium with 10 µM Y-27632 (MedChemExpress) for 24 h to promote cell survival. We maintained all iPSC cultures below passage 50 and confirmed that cultures were negative for mycoplasma before differentiation.

We maintained three human iPSC lines on Matrigel-coated T25 or T75 flasks (Thermo Fisher Scientific) in Essential 8 Flex medium (Gibco) at 37 °C and 5% CO₂. We diluted Matrigel according to the manufacturer-recommended lot-specific dilution, typically 1:100, in ice-cold DMEM/F12 (Gibco) and coated flasks overnight.

We replaced the medium every 2 d and passaged cells after they reached 70-80% confluency. For passaging, we rinsed cultures with HBSS, dissociated cells with Accutase (Gibco) for 3 min at 37 °C, triturated them to a single-cell suspension, diluted the suspension with two volumes of DMEM/F12, and centrifuged at 900 RPM for 5 min. We resuspended the cell pellets in Essential 8 Flex medium and replated the cells at an approximate 1:3 to 1:6 split ratio based on culture surface area.

We cryopreserved iPSCs in CryoKing cryovials at 1 × 10⁶ cells/mL in CELLBANKER® 1 (Zenogen Pharma Co., Ltd., Japan) using controlled-rate freezing at approximately −1 °C/min before transfer to liquid nitrogen.

### OSO generation

We generated OSOs from iPSCs using sequential embryoid body formation, dual-SMAD inhibition, FGF8-mediated olfactory fate specification, and adherent maturation. To form embryoid bodies (EBs), we dissociated iPSCs at 70-80% confluency into single cells using Accutase, quantified viable cells by trypan blue exclusion, and used cultures with >90% viability. We seeded 10,000 cells per well in 200 µL Essential 8 Flex medium containing 10 µM Y-27632 (MedChemExpress) into Nunclon Sphera ultra-low-attachment 96-well plates (Thermo Fisher) and incubated them for 48 hours to allow EB formation.

On Day 2, we induced neural differentiation by dual-SMAD inhibition using 2 µM Dorsomorphin and 10 µM SB431542 (MedChem Express). We additionally supplemented the medium with 10 ng/mL bFGF (MedChem Express) from days 2-4 and continued dorsomorphin and SB431542 (MedChem Express) from days 4-8, with daily medium changes.

On Day 8, we withdrew the dual-SMAD inhibitors and initiated olfactory fate specification by supplementing the medium with 10 ng/mL FGF8. We maintained FGF8 treatment through day 12 with daily medium changes. By day 12, EBs typically reached approximately 400-600 µm in diameter and developed smooth, optically dense peripheries.

From days 30-60, we supplemented the medium with DAPT (5uM) to promote neuronal differentiation. After day 60, we maintained the OSOs in N2+B27-supplemented Neurobasal medium (Gibco). through functional characterization and drug treatment to day 88.

### Brightfield imaging

We acquired brightfield and phase-contrast images using an inverted EVOS XL Core Imaging System (Invitrogen). We calibrated image scale bars before analysis and used the resulting images to document iPSC morphology, embryoid body formation, and OSO development over time.

### Cryosectioning

For cryosectioning, we directly embedded the OSOs in Fisher Healthcare Tissue-Plus O.C.T. Compound (Fisher Scientific, Cat. No. 23-730-571). We then sectioned the embedded OSOs at a thickness of 10 µm using a Leica CM3050 S cryostat (Leica Biosystems) and stored the sections at −80 °C until further use.

### Immunohistochemistry

For immunofluorescence, we washed the OSO sections three times with 1X PBS, blocked them with 10% normal goat serum in 1X PBS for 1 hour at room temperature, and permeabilized them with 0.02% Triton X-100 (Sigma) in 1X PBS for 10 minutes at room temperature. We then incubated the sections overnight (18 h) at 4°C with primary antibodies (**SI Table 1**) diluted in antibody dilution buffer (CST). Following this, we washed the sections three times in 1X PBST and incubated them with Alexa Fluor-conjugated secondary antibodies (ThermoFisher), Alexa Fluor 488 anti-mouse (1:1000) and Alexa Fluor 647 anti-rabbit (1:1000) diluted in antibody dilution buffer (CST) for 1 hour at room temperature. Finally, we washed the sections three times with 1X PBST, counterstained them with the DNA-binding dye DAPI (Invitrogen), and mounted them using an anti-fade mounting medium (Vector Labs). We acquired confocal fluorescence images using an Olympus FV3000RS microscope. The staining panels are shown in **SI Figure 1**.

### snRNA sequencing

We collected OSOs at two and three months, washed them twice with ice-cold DPBS, snap-froze them in liquid nitrogen, and stored them at −80°C before sending them to AmpSeq for nuclei isolation and sequencing. As per AmpSeq protocol nuclei were isolated through detergent-based lysis, collagenase II digestion, and filtration through a 20 µm strainer. Isolated nuclei were stained with DAPI and counted using a fluorescence microscope. Libraries were prepared using the M20 Genomics VITApilote high-throughput single-cell transcriptome kit on the VITAcruizer DP400 platform, following the manufacturer’s protocol. RNA was reverse transcribed within intact nuclei, and the resulting cDNA was barcoded using cell-specific barcodes and unique molecular identifiers (UMIs), amplified, and purified. Libraries were sequenced on an Element Biosciences AVITI platform using 76 bp paired-end reads. Sequencing generated 44,048,864 read pairs and 2,211 nuclei for the two-month sample, and 38,501,743 read pairs and 2,004 nuclei for the three-month sample. Raw sequencing data were processed using VITAseer (VITAbasic v1.5) for adapter trimming, quality filtering, and barcode processing. Cleaned reads were aligned to the human reference genome (GRCh38) using STARsolo (STAR v2.7.10b), with gene expression quantified using both exonic and intronic reads. Nuclei were identified using EmptyDrops for the two-month sample and UMI-based barcode selection for the three-month sample.

### snRNA sequencing data processing

We analyzed the two and three month points separately in Scanpy. We removed low-quality nuclei, retaining those with at least 500 UMIs and 300 genes at two months, or 400 UMIs and 250 genes at three months, and with mitochondrial counts below 10%. This left 1,720 nuclei at two months and 1,565 at three months. We show quality control of the retained nuclei, including UMIs per nucleus, genes per nucleus, and mitochondrial content in **SI Figure 2**. Retained nuclei had a median of 2,325 UMIs and 1,586 genes at two months (median mitochondrial content 4.0%) and 1,135 UMIs and 905 genes at three months (median mitochondrial content 4.6%). We removed genes detected in fewer than three nuclei, normalized counts to 10,000 per nucleus, and log-transformed the data. We selected the 2,000 most highly variable genes, scaled expression to a maximum of 10, and computed 50 principal components. We built the neighborhood graph from the first 30 components and embedded the nuclei with UMAP. We clustered nuclei with the Leiden algorithm at resolution 1.3 for the two-month sample, which gave 14 clusters, and at resolution 1.5 for the three-month sample, which gave 12 clusters. We identified cluster markers with a Wilcoxon rank-sum test and assigned cell-type labels by comparing those markers with established olfactory epithelial and olfactory bulb signatures.

### Pseudotime analysis

We performed pseudotime analysis separately at each time point to follow olfactory maturation. We built a diffusion map with 15 components, set the root to the nucleus with the lowest GNAL expression, and inferred pseudotime with 10 diffusion components. For each marker, we divided nuclei into 100 pseudotime bins, calculated the mean expression in each bin, and smoothed the curve with a Gaussian filter. We plotted mature sensory-neuron genes (GNAL, GNG8, RTP1, CNGA2), periglomerular dopaminergic genes (DDC, SLC6A3), granule and GABAergic interneuron genes (GAD1, GAD2, DLX1, DLX2, DLX5, DLX6), and ensheathing-glia genes (S100B, PLP1, SOX10, ERBB3) at both time points, and examined olfactory-receptor candidates in the same way in the three month OSOs.

### Human fetal data cross-comparison

We compared two month and three month OSO snRNA sequencing datasets with the processed human fetal nasal atlas reported by Mbouamboua et al. (2026).^[31]^ We restricted the analysis to 10,889 genes shared across all datasets and integrated 45,848 nuclei using Seurat and Harmony. We performed Louvain clustering on the Harmony-corrected representation and generated a shared UMAP embedding. We assigned cluster names using fetal cell-type annotations from the reference atlas.

We quantified fetal and OSO representation within the shared clusters and calculated the distribution of nuclei across major neural and olfactory populations. For directional concordance analysis, we defined fetal transcriptional signatures for neural clusters containing both fetal and OSO nuclei and determined whether corresponding gene-expression differences within two- and three-month OSOs changed in the same direction as the fetal signatures. We show detailed preprocessing, clustering, composition, and concordance procedures, and the cluster summary in **SI Table 2**, **Supporting Information**.

### Electrophysiology

We assessed spontaneous neuronal network activity in OSOs with the MED64 Presto multielectrode array (MEA) system (AutoMate Scientific) on 48 well plates with eight electrodes each. We coated the MEA plate first with laminin and then poly-D-lysine and incubated overnight. <u>Maturation study</u>: To track longitudinal functional maturation, we recorded electrophysiological activity for 5 min at weekly intervals from day 60 (week 0) through day 88 (week 4). We followed the same OSOs across all time points (n = 24). From each recording, we quantified network burst duration, network burst rate, spikes per network burst, spike frequency, normalized network burst-duration interquartile range (IQR), and interspike interval to characterize how spontaneous and coordinated neuronal activity changed during maturation.

### Drug dosing study

We treated three-month OSOs with Dasatinib (1 µM), Trametinib (1 µM), or Selumetinib (10 µM), obtained from MedChemExpress. We selected these concentrations based on previously published pharmacological studies. Dasatinib has been investigated in clinical studies and evaluated at 1 µM in cancer cell lines to assess its inhibitory effects on Src/Abl kinase signaling.^[48,49]^ Trametinib has been evaluated at 1 µM in vitro to investigate MEK inhibition and downstream signaling responses.^[50]^ Selumetinib was administered at 10 µM, a concentration previously examined in kinase selectivity assays.^[51]^

To provide reference conditions for assessing neuronal excitability, we additionally treated OSOs with potassium chloride (KCl, 25 µM) as a depolarizing reference and gamma-aminobutyric acid (GABA, 25 µM) as an inhibitory reference. The kinase inhibitors were prepared in DMSO, and DMSO-treated OSOs served as vehicle controls. The final DMSO concentration was maintained at ≤0.1% across treatment conditions.

Immediately before treatment, we recorded baseline network activity using the MED64 Presto multielectrode array system. Following drug administration, we recorded electrophysiological activity every 2 h for 24 h. We quantified average spike frequency, average interspike interval, spikes per network burst, and network burst rate and compared these measurements across treatment and vehicle-control conditions.

## Statistical analysis

We performed all statistical analyses in GraphPad Prism (version 11.2.0) and present data as mean ± standard error of the mean (SEM). We analyzed each electrophysiological parameter separately with Dunnett’s multiple comparisons test. In the maturation study, Dunnett’s test compared weeks 1 through 4 with week 0. In the pharmacological study, it compared KCl, GABA, Dasatinib, Trametinib, and Selumetinib with DMSO vehicle. Dunnett’s test controls the family-wise error rate across comparisons against a shared reference. For each comparison, we reported the multiplicity-adjusted p value together with the mean difference and its 95% confidence interval. All tests were two-sided, and we defined statistical significance as an adjusted p < 0.05 (*p < 0.05, **p < 0.01, ***p < 0.001, ****p < 0.0001).

## Supplementary List

Gene lists for the clusters in **Figure 2** and the values plotted in the pseudotime panels are provided as supplementary CSV files. Fig2B_olf_two month_genelist.csv and Fig2C_olf_three month_genelist.csv list the cluster genes for the two-month and three-month OSOs. Fig2D-E_plotted_pseudotime_control_markers_OR_Candidates.csv contains the plotted pseudotime values for the three-month transduction markers and odorant-receptor candidates (Figure 2D,E). Fig3A-D_plotted_pseudotime_OSN.csv contains the plotted values for the sensory-neuron genes (**Figure 3A-D**). Fig3E-F_plotted_pseudotime_DA_GABA.csv and Fig3G-L_plotted_pseudotime_DA_GABA.csv contain the plotted values for the dopaminergic and GABAergic genes (**Figure 3E-L**). Fig3M-P_plotted_pseudotime_Glia.csv contains the plotted values for the ensheathing-glia genes (**Figure 3M-P**). Fig5_panel_b_Human_fetal_nasal_atlas_crosscomparison_cluster_summary.xlsx lists all 24 clusters of the shared map from the Harmony integration, with the main fetal cell types and the number of fetal and organoid nuclei in each (**Fig. 5B**). Fig5_panel_c_fetal_vs_organoid_share_in_each_cluster.csv gives the number and percentage of fetal and organoid nuclei in each cluster shown in Fig. 5C. Fig5_panel_di_concordance_cluster02_neurons.csv, Fig5_panel_dii_concordance_cluster10_neural_progenitors.csv, Fig5_panel_diii_concordance_cluster16_immature_OSNs.csv and Fig5_panel_div_concordance_cluster23_NOS1_neurons.csv give, for each fetal signature gene, the fetal and organoid values plotted in Fig. 5**D**(**i**)-(**iv**) for neurons, neural progenitors, immature olfactory sensory neurons and NOS1+ neurons. Fig5_panel_e_where_each_sample’s_nuclei_fall.csv gives the number and percentage of nuclei in each cluster group for each organoid age and for the fetal neural and olfactory epithelium nuclei (**Fig. 5E**).

## Acknowledgements

The manuscript’s grammar was improved using ChatGPT (OpenAI, 2026) and Grammarly, while maintaining all scientific content and conclusions by the authors. Schematics in Figures 1, 2, 4, and 5 were created using BioRender (Created in BioRender. Kathuria, A. (2026). https://BioRender.com/8rqyz3g).

## Author Contribution

V.S.: Formal analysis (lead); methodology (co-lead); Conceptualization (equal); data curation (equal); investigation (equal); visualization (lead); writing – original draft (lead); writing – review and editing (equal). A.P.: Conceptualization (equal); data curation (equal); formal analysis (equal); investigation (equal); methodology (co-lead); writing – review and editing (equal). K.J.: Investigation (equal); methodology (equal); writing – review and editing (equal). O.S.: Data curation (equal); formal analysis (equal); investigation (equal); methodology (equal); writing – review and editing (equal). N.L.: Investigation (equal); methodology (equal); writing – review and editing (equal). A.K.S.: Investigation (equal); methodology (equal); writing – review and editing (equal). N.Li.: Data curation (equal); formal analysis (equal); investigation (equal); methodology (equal); writing – review and editing (equal). X.Z.: Formal analysis (equal); investigation (equal); methodology (equal); supervision (equal); writing – review and editing (equal). V.M.: Formal analysis (equal); investigation (equal); methodology (equal); supervision (equal); writing – review and editing (equal). A.K.: Conceptualization (lead); data curation (equal); formal analysis (equal); funding acquisition (lead); investigation (lead); methodology (equal); project administration (lead); supervision (lead); writing – review and editing (equal).

## Data Availability Statement

The authors declare that the data that support the findings of this study are available from the corresponding author upon request.

## Supporting Information

Supporting Information is available from the Wiley Online Library or from the author. Human iPSC-derived olfactory sensory organoids are specified with FGF8 and matured for three months. snRNA sequencing and pseudotime analysis resolve sensory, interneuron, and glial programs that align with a fetal nasal atlas. Network activity recorded on multielectrode arrays matures over time and changes in a compound-specific way after Dasatinib, Trametinib, or Selumetinib.

